# Data-driven polymer model for mechanistic exploration of diploid genome organization

**DOI:** 10.1101/2020.02.27.968735

**Authors:** Yifeng Qi, Alejandro Reyes, Sarah E. Johnstone, Martin J. Aryee, Bradley E. Bernstein, Bin Zhang

## Abstract

Chromosomes are positioned non-randomly inside the nucleus to coordinate with their transcriptional activity. The molecular mechanisms that dictate the global genome organization and the nuclear localization of individual chromosomes are not fully understood. We introduce a polymer model to study the organization of the diploid human genome: it is data-driven as all parameters can be derived from Hi-C data; it is also a mechanistic model since the energy function is explicitly written out based on a few biologically motivated hypotheses. These two features distinguish the model from existing approaches and make it useful both for reconstructing genome structures and for exploring the principles of genome organization. We carried out extensive validations to show that simulated genome structures reproduce a wide variety of experimental measurements, including chromosome radial positions and spatial distances between homologous pairs. Detailed mechanistic investigations support the importance of both specific inter-chromosomal interactions and centromere clustering for chromosome positioning. We anticipate the polymer model, when combined with Hi-C experiments, to be a powerful tool for investigating large scale rearrangements in genome structure upon cell differentiation and tumor progression.

## Introduction

Three-dimensional genome organization is essential for DNA templated processes, including transcription, DNA replication, and repair.^1,2^ Advances in chromosome-conformation-capture (Hi-C) and related methods have greatly improved its high-resolution characterization,^3^ and have led to the discovery of chromatin loops,^4^ topologically associating domains (TADs),^5,6^ and compartments^7^ at various scales. Much progress has been made towards unraveling the molecular mechanism of loop and TAD formation as well.^8^ In particular, the loop extrusion model that hypothesizes a processive movement of cohesin molecules along the DNA^9,10^ helps to explain the co-localization of CCCTC-binding factor (CTCF) and cohesin at TAD boundaries. Numerous predictions of the extrusion model have been validated with perturbative Hi-C and single-cell super-resolution imaging experiments.^11–14^ Most recently, it has been proposed that loop extrusion and compartmentalization are both crucial for the formation of TADs.^15,16^

Though much is known about the structure and folding of individual chromosomes, questions regarding the genome organization at a global scale remain outstanding.^17–19^ In particular, imaging studies have long demonstrated that chromosome nuclear localizations are non-random and highly correlated with transcriptional activity.^20^ What drives the relative positioning of individual chromosomes is not well understood, however. Such questions are challenging to address experimentally since high-throughput sequencing-based techniques measure contact frequencies via proximity ligation and do not directly report 3D positions. In addition, due to its inherent limitation in detecting rare contacts that are separated farther apart in space,^21,22^ the usefulness of Hi-C for characterizing inter-chromosomal interactions remains to be shown. On the other hand, imaging-based techniques, though ideally suited for spatial measurements, are often low throughput and can face challenges for mechanistic explorations that require large data set collection for statistical significance.

As a complementary approach, polymer simulations can be useful for studying genome organization as well.^23^ In particular, numerous groups have developed computational techniques to reconstruct high-resolution chromosome structures as polymer models that recapitulate Hi-C data.^24–28^ These studies have provided great details into the organization of both interphase and metaphase chromosomes. However, it is often difficult to address questions regarding why chromosomes adopt certain structures with these purely data-driven approaches. Alternatively, great insight can be gained by engineering specific mechanisms of genome organization into polymer simulations and evaluating the quality of resulting chromosome structures. Such hypothesis-driven approaches have demonstrated the importance of various biological factors, including protein-mediated loop formation,^29–33^ phase separation and compartmentalization,^15,34^ and non-equilibrium dynamics^35–37^ in genome organization. Due to the presence of free parameters whose values cannot be determined straightforwardly, generalizing these approaches to evaluate the relative significance of various mechanisms and identify a minimum set of hypotheses that are both necessary and sufficient for modeling genome organization is difficult.

Coupling the data- and hypothesis-driven approaches together can potentially produce a powerful strategy for modeling genome organization.^38–40^ This strategy ensures the biological relevance of simulated genome structures since all model parameters will be derived from Hi-C data. In the meantime, it will be well suited for mechanistic investigation as the polymer model’s energy function will be designed explicitly from biological factors that are known to contribute to genome organization. Recently, we applied this strategy to model individual chromosomes at a 5kb resolution by parameterizing a chromatin-state based energy function from Hi-C data.^38^ Starting from one-dimensional genomics and epigenomics data that are available for hundreds of cell types, the model provides a high-resolution characterization of various chromatin structural motifs that is in quantitative agreement with Hi-C and super-resolution microscopy measurements. In addition, it uncovers numerous chromatin features that contribute to cell-type specific enhancer-promoter contacts.

Here, we generalize the data-driven mechanistic-modeling approach to study the organization of the diploid genome. We model chromosomes at 1Mb resolution and explicitly consider forces that drive *A*/*B* compartmentalization, centromere clustering, and *X* chromosome inactivation. Parameters that evaluate the strength of these interaction forces were derived using only haploid-specific Hi-C data. We find that, despite its simplicity, the polymer model succeeds in quantitatively reproducing Hi-C data, including intra- and inter-chromosomal contacts. Simulated genome structures are also in qualitative agreement with a wide variety of imaging studies, including the preferential interior localization of active chromosomes, the radial positioning of individual chromosomes, the condensation of the inactive *X*-chromosome, centromere clustering, etc. In addition, homolog-specific structural features are in good agreement with models built from single-cell diploid Hi-C data as well. Detailed mechanistic investigations suggest that the nuclear localization of individual chromosomes is not fully driven by the phase separation of *A*/*B* compartments; both centromere clustering and specific inter-chromosomal interactions contribute significantly to chromosome positioning. Our study, therefore, establishes a modeling framework for efficient and accurate reconstruction of homolog-resolved whole-genome organization from haploid Hi-C data.

## Results

### A data-driven mechanistic model for the diploid human genome

We introduce a polymer model to study the 3D organization of the diploid human genome in interphase. As shown in Fig. 1, each one of the 46 chromosomes is modeled as a string of beads that are 1Mb in length. Though fine structural details such as chromatin loops are inevitably lost at this resolution, global features of genome organization, including territory formation and compartmentalization, can be readily recognized and captured.^7^ We further differentiate the beads as either compartment *A* or *B* based on the corresponding interaction patterns detected in Hi-C contact maps. Since most Hi-C data are processed at the haploid level, identical compartment profiles were used for the two homologous copies of each chromosome. Prior studies with diploid-specific Hi-C data indeed support the validity of this approximation,^4,41^ and the rare single-nucleotide polymorphisms (SNPs) or imprinted loci do not contribute significantly to the megabase level organization considered here.

**Figure 1:**
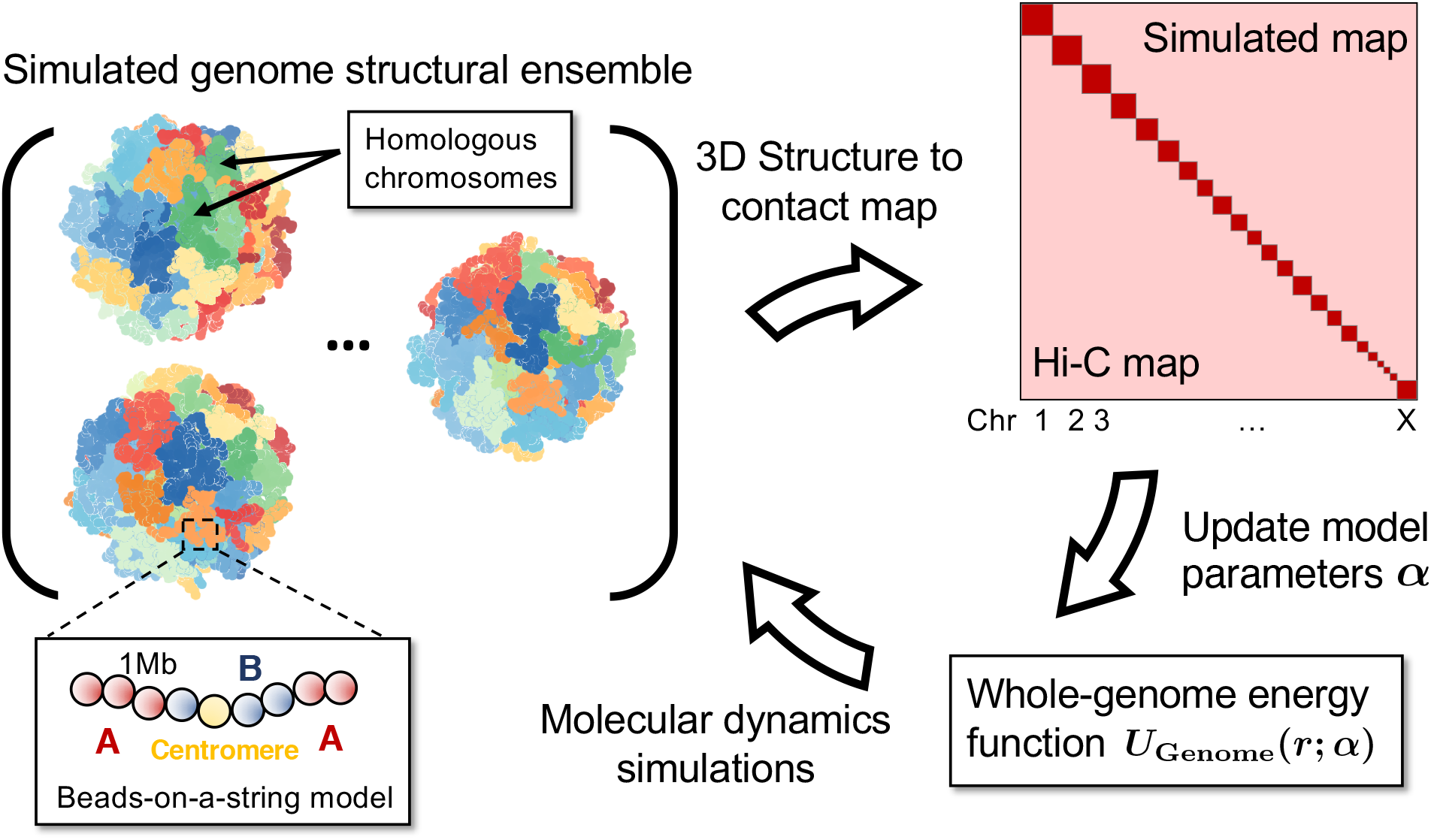
Overview of the iterative algorithm for parameterizing the polymer model of diploid genome organization. Each one of the 46 chromosomes is explicitly represented at the 1Mb resolution as a string of beads, which are further distinguished as compartment *A*, *B* or *C*. Given a set of parameters that measures the interaction strength among chromosomes, we use molecular dynamics simulations to collect an ensemble of genome structures. By converting these structures into a haploid contact map, polymer simulations can then be directly compared against Hi-C data. The difference between the two will provide further guidance on updating the parameters.

The block copolymer model based on *A*/*B* compartments outlined above is too simplistic to capture the full complexity of genome organization, however.^42^ We added the following modifications to better describe inter-chromosomal interactions. First, we defined a new type *C* to recognize centromeric regions and distinguish them from the *A*/*B* compartments. Separating out the centromeric regions is necessary to capture the specific interactions among them that can significantly impact the localization of numerous chromosomes.^25,43,44^ Second, we modeled intra- and inter-chromosomal interactions with two sets of parameters to account for the presence of distinct driving forces for chromosome contacts at different length scales. In particular, numerous dynamical processes powered by processive motors that track along the DNA molecule, including loop extrusion by cohesin molecules^9,10^ and transcription by RNA polymerase,^45^ only contribute to the organization of individual chromosomes. On the other hand, contacts between chromosomes can arise independently from these processes as a result of their mutual interaction with the same nuclear landmark such as the envelope and the nucleolus.^8,46^ Third, to capture the structural difference between the two *X* chromosomes, we applied additional intra-chromosomal interactions for the inactive copy inspired by its association with lncRNA molecules that promote chromatin condensation.^47,48^ Lastly, recognizing that the polymer model built upon *A*/*B* compartments cannot resolve any DNA sequence specific effects, we introduced correction terms to each individual pair of chromosomes to further improve the model’s accuracy.

The potential energy function of the whole genome model detailed above can be written as the following

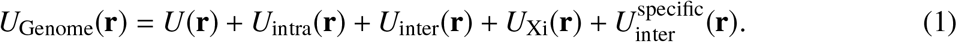

*U*(**r**) is a generic polymer potential applied to all chromosomes to define the topology of the polymer, the excluded volume effect among genomic loci and the volume fraction of the DNA inside the nucleus. *U*_intra_(**r**) = Σ_*I*_ Σ_*i, j*∈*I*_ *u*(*r*_*i j*_, *T*_*i*_, *T*_*j*_) describes the type-specific intra-chromosomal potential, with *I* indexing over different chromosomes. The strength of the interaction energy *u* between a pair of genomic loci *i* and *j* depends both on their distance *r*_*i j*_ and the chromatin type *T*, which can be *A*, *B* or *C*. *U*_inter_(**r**) = Σ_*I,J*_ Σ_*i*∈*I*, *j*∈*J*_ *v*(*r*_*i j*_, *T*_*i*_, *T*_*j*_) is similarly defined but for interactions between genomic loci from different chromosomes. The weakly attractive potential *U*_Xi_(**r**) = Σ_*i, j*∈Xi_ *w*(*r*_*i j*_) is only applied to the inactive *X*-chromosome. 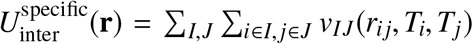 defines a specific interaction potential for every pair of heterologous chromosomes. Detailed expressions for the different energy terms are provided in the *Methods Section*. We note that *U*_Genome_(**r**) is designed to be general so that it can capture the diverse set of mechanisms of genome organization. It does not enforce any of the hypotheses mentioned above, however. As shown below, the relative strength of different energy terms will be derived from Hi-C data. If the experimental data does not support a particular hypothesis, the strength of the corresponding energy will be automatically set to zero.

While the polymer model is motivated to recapitulate various biological mechanisms of genome organization, it is also data-driven and all of its parameters can be determined from haploid Hi-C data with an iterative optimization algorithm.^27,38^ Model parameterization is possible because mathematical expressions for the various energy terms in *U*_Genome_(**r**) were designed such that their ensemble averages can be mapped onto combinations of contact frequencies measured in Hi-C. As shown in Fig. 1, we first perform molecular dynamics simulations to collect an ensemble of genome structures. By averaging over homologous chromosomes, we then convert the simulated structures into a haploid contact map as similarly done in Hi-C data processing. From the simulated contact map, constraints that correspond to the different energy terms can be determined and compared with the counterparts estimated using Hi-C data. For constraints involving centromeric regions, we approximated the experimental values using contact frequencies estimated from peri-centromeric regions due to the lack of Hi-C data. Finally, model parameters can be updated based on the difference between simulated and experimental constraints. With the newly updated parameters, one can restart the three steps if necessary to further improve the agreement between simulation and experiment. More details on parameter optimization can be found in the supporting information (SI).

### Simulated contact map reproduces experimental Hi-C data

We applied the optimization algorithm to derive a whole genome model for GM12878 cells. Using the parameterized energy function, we simulated the ensemble of genome structures and determined the contact probability map. As shown in Fig. 2a, the contact map supports the formation of chromosome territories and intra-chromosomal interactions are much stronger than the inter-chromosomal ones. The mean contact probabilities within each chromosome (Fig. 2b) and the contact probability as a function of genomic separation averaged over all chromosomes (Fig. 2c) both match well with the corresponding experimental values. Further zooming in on individual chromosomes reveals that large-scale compartmentalization is reproduced satisfactorily (Fig. 2d), though some fine features seen in Hi-C along the diagonal are evidently missing in the simulated contact map. Capturing such detailed interaction patterns potentially requires the use of sub-compartment types or chromatin states to resolve the varying degree of active (*A*) and inactive (*B*) chromatin.^4,38^ However, we found that these finer structures were not necessary for an accurate modeling of chromosome positioning and inter-chromosomal interactions, indicating that sub-compartments might only contribute to local chromatin folding rather than global genome organization.

**Figure 2:**
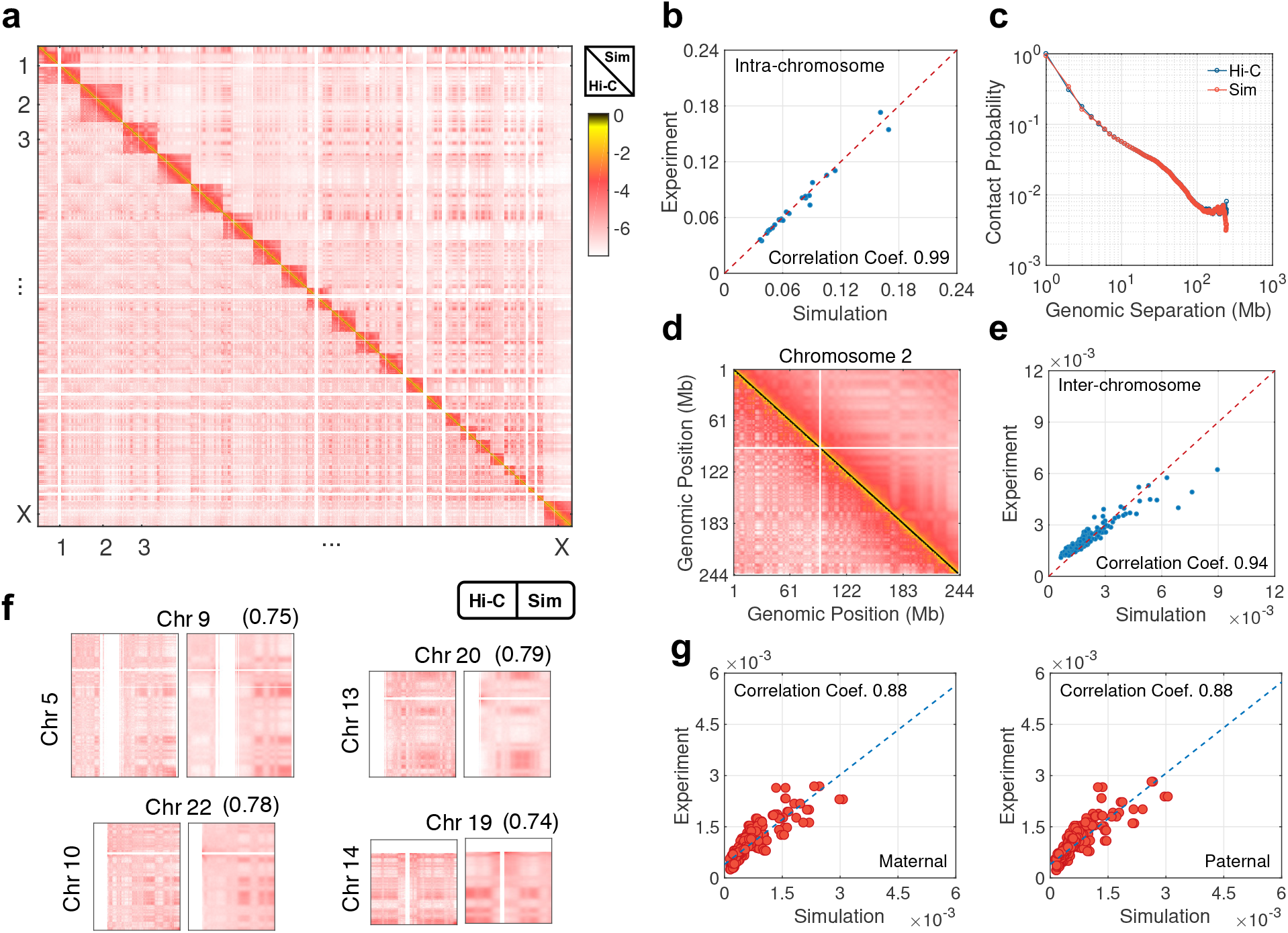
Polymer simulations succeeds in reproducing intra- and inter-chromosomal contacts measured in Hi-C experiments.^4^. (a, b, c, e) Comparison between simulated and experimental, whole genome contact map at the 1Mb resolution (a), average intra-chromosomal contact probabilities (b), the contact probability as a function of genomic separation averaged over all chromosomes (c), and average inter-chromosomal contact probabilities (e). (d) A zoomed-in view of the whole genome contact map shown in part a for chromosome 2, with simulated and experimental results shown in the upper and lower triangle, respectively. (f) Zoomed-in views of representative interchromosomal contacts. Chromosomes IDs are provided on the side and the Pearson correlation coefficients between simulated and experimental inter-chromosomal contact matrices are shown in parentheses. (g) Comparison between simulated and experimental average inter-chromosomal contacts formed with maternal (*Left*) or paternal (*Right*) chromosomes.

Next, we examined the quality of simulated inter-chromosomal contacts. As shown in Fig. 2e, the average contact probabilities between pairs of chromosomes agree well with the experimental values. Close examination of inter-chromosomal contact blocks, with examples provided in Fig. 2f, suggests that the model succeeds in recapitulating specific interaction patterns as well. Pearson correlation coefficient between these blocks is around 0.65 when averaged over all chromosome pairs (Fig. S1). This correlation coefficient is comparable to the one between experimental replicates (Pearson correlation coefficient = 0.78).

Homologous autosomes in our polymer model share identical compartment profiles and interact with the rest of the genome in the same way. To validate the accuracy of this assumption, we reprocessed the Hi-C data^4^ to build a diploid contact map using allele-specific SNPs information (see *Method Section*). Since contacts with both DNA strands containing at least one SNP is extremely rare, only the parental origin for one of the loci contributing to the contact is resolved. The inter-chromosomal contacts are now, therefore, split into two sets that correspond to contacts originating from maternal or paternal chromosomes. As shown in Fig. S2a, these two data sets are well correlated with each other, thus providing strong support for our approximate treatment of homologous chromosomes.

In Fig. 2g, we further compare the simulated allele-specific average contact probabilities between pairs of chromosomes with those estimated from diploid Hi-C contacts. Since the polymer model does not distinguish the two homologs, we randomly assigned one of them as the maternal copy and the other one as the paternal copy for the convenience of data processing. The simulated results are in good agreement with the experimental values, with a Pearson correlation coefficient of 0.88. As expected, flipping the maternal and paternal assignment in the polymer model does not lead to noticeable changes in the resulting contact probabilities (Figs. S2b and S2c).

### Simulated polymer structures recapitulate 3D genome organization

Encouraged by its success in reproducing Hi-C data, we now analyze the structural details of the polymer model. A snapshot for the simulated genome organization is provided in Fig. 3a, with heterologous chromosomes marked in distinct colors for better visualization. The model clearly reproduces the formation of spatially separated territories for individual chromosomes.^49–51^ In addition, it also predicts a large difference in terms of morphology for the two *X* chromosomes, with the inactive one being more compact and adopting a more spherical shape (also see Fig. 3c and Fig. S3).

**Figure 3:**
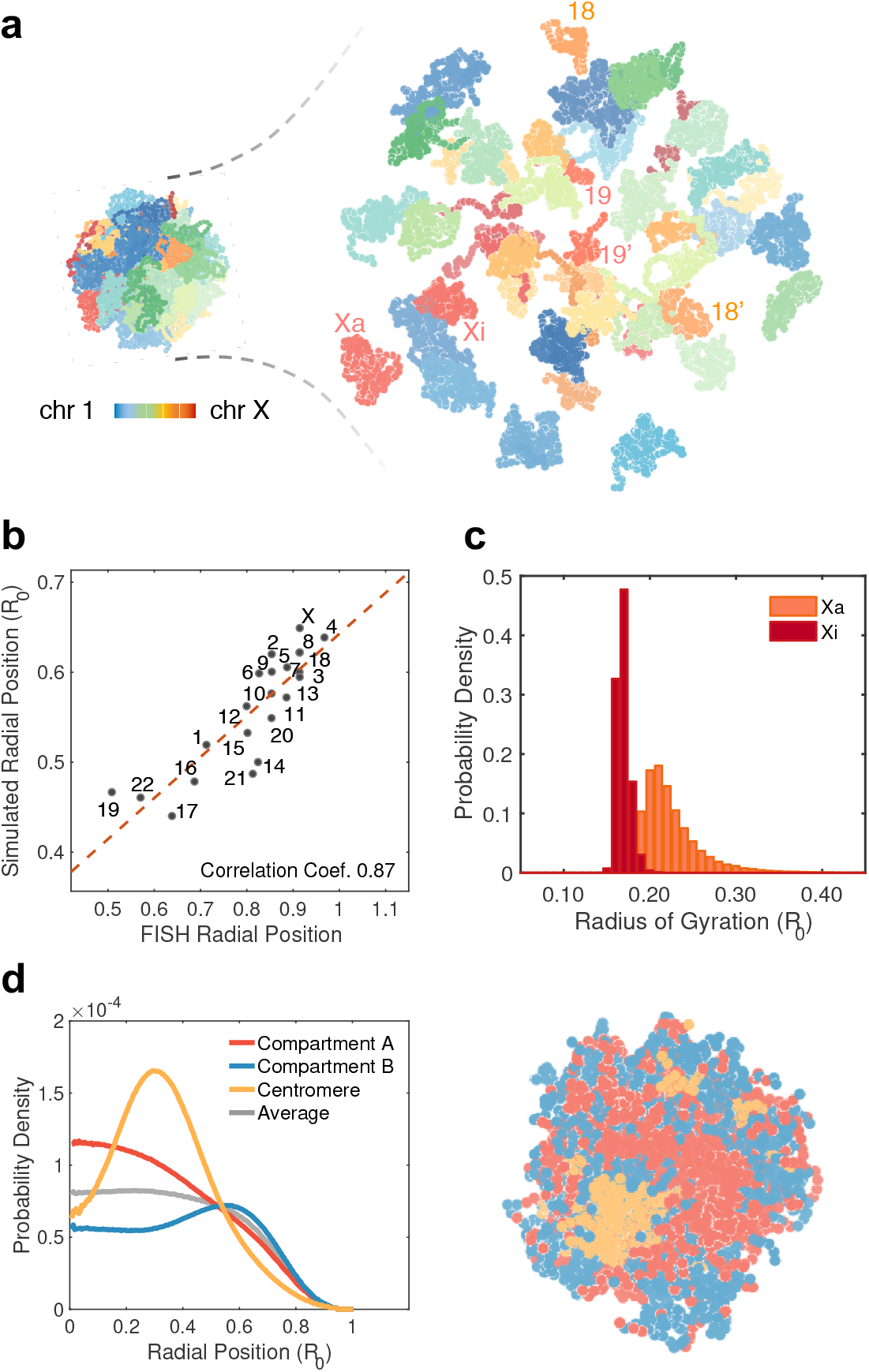
Polymer simulations reproduce large scale genome organization. (a) Example genome structure predicted by the polymer model with chromosomes shown in different colors. A blow-up version is shown on the side. (b) Comparison between simulated and experimental^20^ average radial position of individual chromosomes. (c) Probability distribution of the radius of gyration for the active (orange) and inactive (red) *X*-chromosome. (d) Radial distribution of various compartment types as a function of the distance from the nuclear center. An average distribution of all genomic loci is shown in grey for reference. A cross section of a genome structure is shown on the side using the same coloring scheme. *R*_*o*_ is the radius of the nucleus used in polymer simulations.

Prior studies have suggested that the spatial organization of the genome is nonrandom and chromosomes adopt well-defined radial positions.^20,52^ Fig. 3a indicates that the model succeeds in recapitulating the well-known observation that the gene-rich chromosome 19 is closer to the nuclear interior than the gene-poor chromosome 18.^20,50,53,54^ For a more systematic comparison, we determined the 3D localization of each chromosome as the average distance from its center of mass to the nuclear origin. As shown in Fig. 3b, the polymer model quantitatively reproduces the data from chromosome painting experiments performed by Bickmore and coworkers.^20^ Additional analyses further revealed that simulated chromosome positions are strongly correlated with gene density, Lamina-associated domain density, and chromosome compactness (Fig. S4).

In addition to chromosome-specific behavior, we examined the spatial distribution of various compartment types as well. As shown in Fig. 3d, the genome exhibits a clear phase separation, with *A* compartments (euchromatin) localizing at the nuclear interior, while *B* compartments (heterochromatin) remain close to the nuclear envelope. This separation is also evident from the cross-section of a representative structure shown on the right. Fig. 3d further reveals that centromeric regions do not mix with *A*/*B* compartments, but form spatial clusters that localize more or less in between the two compartments. These observations are indeed consistent with results from fluorescence in situ hybridization (FISH) experiments that probe the 3D localization of genomic regions corresponding to distinct compartment types.^18,25,54–61^

To evaluate the quality of homolog-specific predictions, we compared the simulated genome structures with those reconstructed from single-cell diploid Hi-C data.^62^ As demonstrated by the pair-wise distance maps between all 46 chromosomes (Fig. 4a), the two structural models bear a resemblance. They both support that, consistent with their interior nuclear localization, chromosomes shorter in sequence length (chr15-chr22) reside in spatial proximity. Chromosome 18, however, appears to be an outlier of this trend. Given the difference in the experimental data and computational algorithms used for their reconstruction, the apparent agreement between the two sets of structures with a correlation coefficient of ~0.6 is significant. In addition, homologous chromosomes, in general, are found to be far apart from each other, and the distances between shorter homologous chromosomes are smaller compared to that of the longer chromosomes (Fig. 4b). These distances are comparable or slightly smaller than the ones between heterologous chromosomes (Fig. 4c).

**Figure 4:**
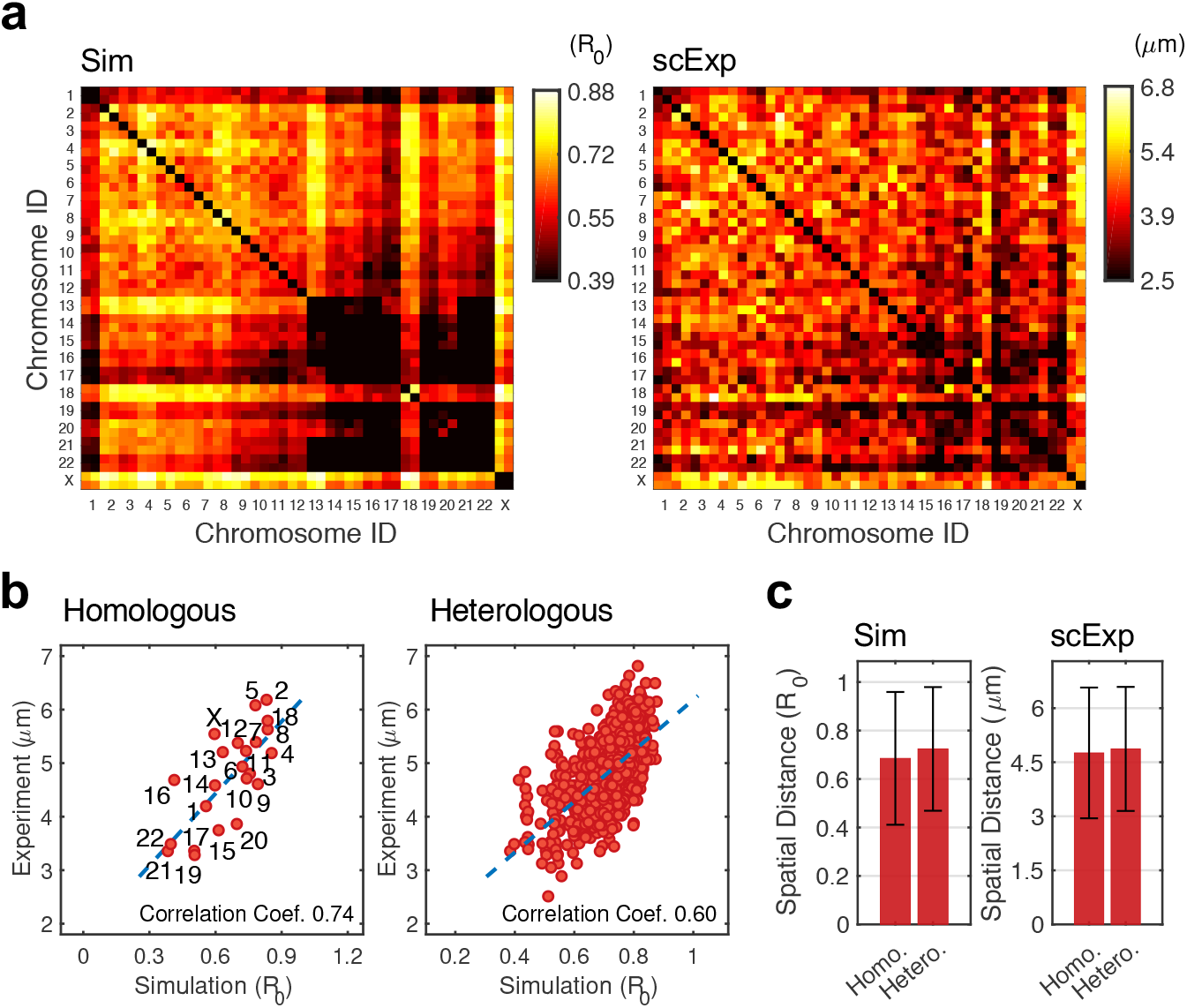
Homolog-resolved genome organization from polymer simulations agrees well with structures reconstructed from single-cell diploid Hi-C data^62^. (a) Average pair-wise distance maps between all 46 chromosomes determined from polymer simulations (*Left*) and from single-cell structures (*Right*). (b) Spatial distances between pairs of homologous (*Left*) or heterologous (*Right*) chromosomes determined from polymer simulations and single-cell structures are strongly correlated. (c) Average distance between homologous and heterologous chromosomes from polymer simulations (*Left*) and single-cell structures (*Right*). Error bars correspond to the variance over all chromosomes. *R*_*o*_ is the radius of the nucleus used in polymer simulations.

The polymer model, therefore, succeeds in reproducing a wide range of qualitative and quantitative imaging and single-cell results. It is important to emphasize that only Hi-C data were used for the model parameterization and 3D genome structures are in fact *de novo* predictions of the model. The agreement between simulated structures and prior independent studies provides strong support that the hypotheses introduced in designing the model energy function are sufficient for a faithful description of the whole genome organization.

### Both specific and non-specific inter-chromosomal interactions contribute to genome organization

A significant advantage of the polymer model is its simplicity and the clear biological meaning of the different terms in the potential energy function. Therefore, perturbations can be performed to selectively remove one or multiple of these terms and evaluate the corresponding impact on genome organization. Such perturbations can help identify mechanisms that are responsible for the model’s success in reproducing large-scale genome organization and serve as effective means to explore the principles of genome organization.

We assumed that intra- and inter-chromosomal interactions differ and must be modeled with two separate potentials. To evaluate the impact of this assumption on genome organization, we built a new polymer model in which a common potential and one single set of parameters was used for both intra- and inter-chromosomal interactions. The iterative algorithm was again employed to ensure that this model reproduces another set of constraints derived from Hi-C data (see SI for details). As shown in Fig. S5a, the block-wise pattern for inter-chromosomal contacts is less evident in the simulated contact maps. Though the two compartments remain largely separated from each other in the simulated structures (Fig. 5a), instead of localizing in the interior, *A* compartments tend to spread over the entire nucleus. Furthermore, simulated radial chromosomal positions are poorly correlated with experimental values, partially caused by the outward movement of the shorter chromosomes. Reproducing the spatial arrangement of chromosomes inside the nucleus, therefore, necessitates a fine tuning of the inter-chromosomal potential. When treated separately from the intra-chromosomal one, the potential between *A* compartments is stronger than that between *B* compartments due to their higher inter-chromosomal contact frequency (Fig. S6b). A stronger interaction potential drives their interior localization to maximize contacts among *A* compartments. If intra- and inter-chromosomal contacts are mixed together, the relative magnitude of the contact frequencies between the two compartments is comparable (Fig. S6c) and so are the interaction energies, resulting in a dramatic change in genome organization.

**Figure 5:**
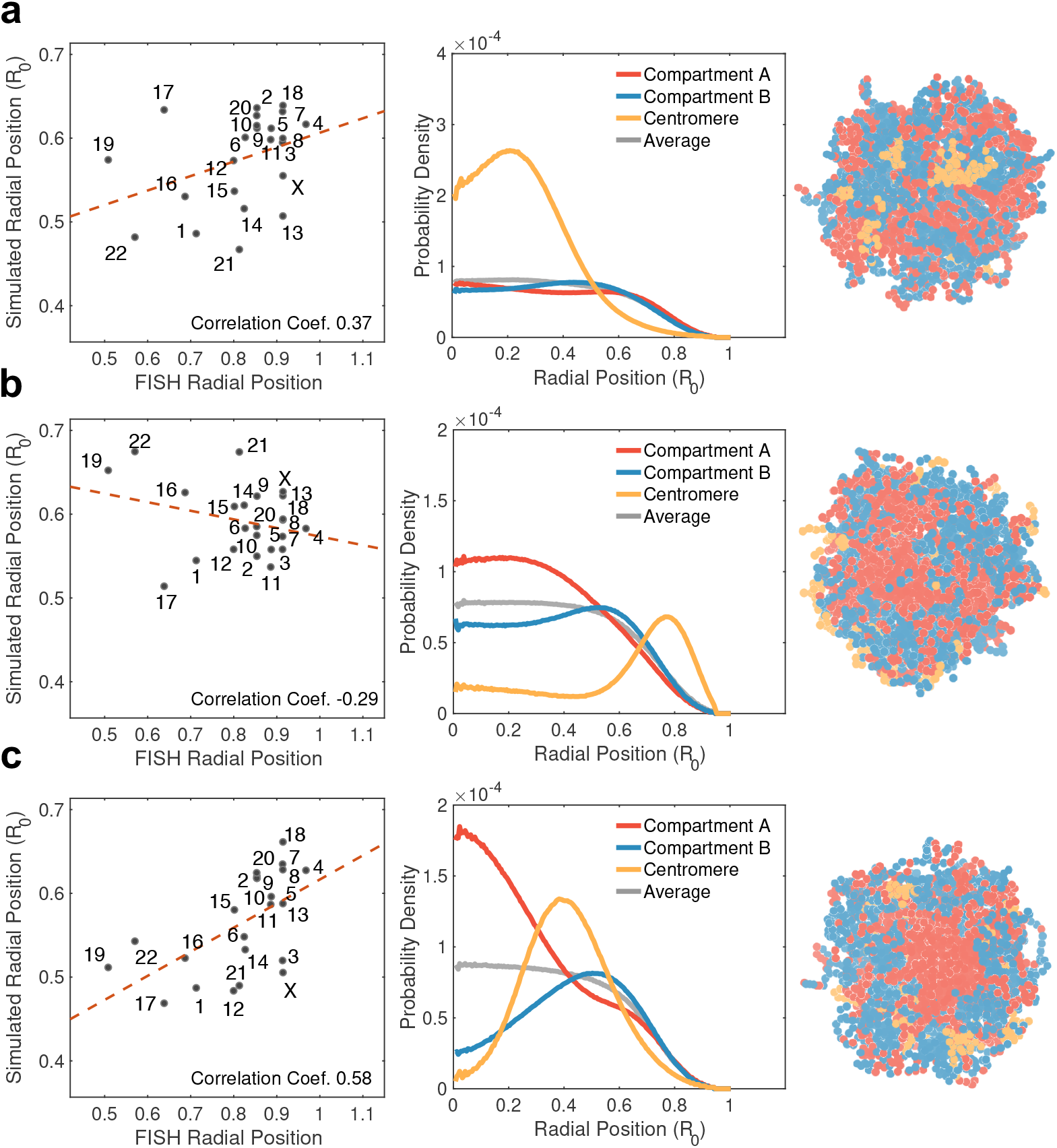
Impact of non-specific and specific inter-chromosomal interactions on global genome organization. Radial positions of individual chromosomes (*Left*), radial distributions of compartment types (*Middle*) and example genome structures (*Right*) are shown for the polymer models that ignores the difference between intra- and inter-chromosomal interactions (a), that removes the interaction among centromeric regions (b), and that abolishes specific inter-chromosomal contacts (c). *R*_*o*_ is the radius of the nucleus used in polymer simulations.

Next, we investigated the role of centromeric regions in organizing the genome with a model that abolishes the specific interactions between them (see SI for details on model parameterization). Comparison between simulated and experimental Hi-C contact maps is provided in Fig. S5b. As shown in Fig. 5b, this perturbation has little impact on the radial distribution of *A*/*B* compartments. However, centromeric regions no longer cluster together in simulated structures but become more scattered at the nuclear periphery. In addition, the radial positions of chromosomes now correlate poorly with the experimental values and many of the shorter chromosomes (chr16, 17, 19, 20 and 21) relocate towards the nuclear periphery. Therefore, *A*/*B* compartmentalization alone does not fully determine the localization of individual chromosomes and clustering among centromeric regions plays a crucial role as well.

Finally, we examined the impact of specific inter-chromosomal contacts on chromosome positioning by parameterizing a polymer model that abolishes the corresponding interaction potential (see SI for details). As shown in Fig. 5c, removing this specific interaction results in suboptimal performance for reproducing chromosome positions, though the overall genome organization appears reasonably preserved (see Fig. S5c as well).

## Discussion

In this work, we presented a polymer model for studying homolog-resolved 3D genome organization. The model is data-driven and all of its parameters can be derived from haploid Hi-C data. Eliminating the uncertainty in model parameters ensures the quantitative accuracy of predicted genome structures. We found that the simulated polymer configurations reproduce a wide range of experimental results independent from the input Hi-C data, including chromosome radial positions, *A*/*B* compartmentalization and inter-homolog distances. These agreements are particularly noteworthy given the model’s simplicity. When designing its energy function, we attempted to reduce the number of parameters based on biologically motivated hypotheses to improve the model’s interpretability. This hypothesis-driven nature makes the model well-suited for mechanistic exploration of genome organization as well.

A key insight from our study is that inter-chromosomal interactions must be separately modeled from intra-chromosomal ones. This separate treatment is partially motivated by the observation that the average intra-chromosomal contact frequency between *A* compartments is weaker than that between *B* compartments, while the opposite trend is observed for inter-chromosomal contacts (Fig. S6). Higher contact frequency leads to a stronger interaction energy between *A* compartments from different chromosomes, resulting in their interior localization. Since *B* compartments within the same chromosome are more condensed compared to *A* compartments due to the formation of heterochromatin, their higher intra-chromosomal interaction energy is indeed reasonable. On the other hand, inter-chromosomal contacts between *B* compartments is mostly driven by their interaction with lamin proteins and co-localization to the nuclear envelope. These “accidental”, indirect interactions are potentially weaker than the specific contacts between *A* compartments from different chromosomes that are of biological significance.^22^

Additionally, we found that the inactive *X*-chromosome is located more interior than the active one (Fig. 6a). This seemingly surprising result is indeed consistent with the observation by Cremer and coworkers.^63^ Using multi-color whole-chromosome painting and quantitative 3D image reconstruction, they found that a significant portion of both chromosomes are located near the nuclear rim while the inactive one extends further into the nuclear center. The authors postulated that its more interior localization could potentially be driven by the specific interaction between the inactive *X* chromosome and the perinucleolus compartment. Alternatively, our polymer simulations suggest that the more compacted conformation of the inactive *X*-chromosome could contribute to its interior localization as well. First, we note that, as shown in Fig. 6b, there is a favorable and comparable energetic gain for both *X* chromosomes to move inward due to their increased contact with the rest of the genome. This energetic driving force, however, is inevitably counterbalanced by the penalty arising from a loss of the configurational entropy. As chromosomes move inward, they become more compact and the set of configurations they can adopt becomes more limited. The entropic penalty would be more significant for the active chromosome given its more expanded conformations, preventing it from occupying more interior locations. We further verified this entropy-driven mechanism by examining the dependence of the radial positions of the two *X* chromosomes on temperature. At higher temperature, the impact of configurational entropy is expected to be stronger and the mechanism would predict a larger difference in the radial positions. Our simulations shown in Fig. 6c are in excellent agreement with such a prediction.

**Figure 6:**
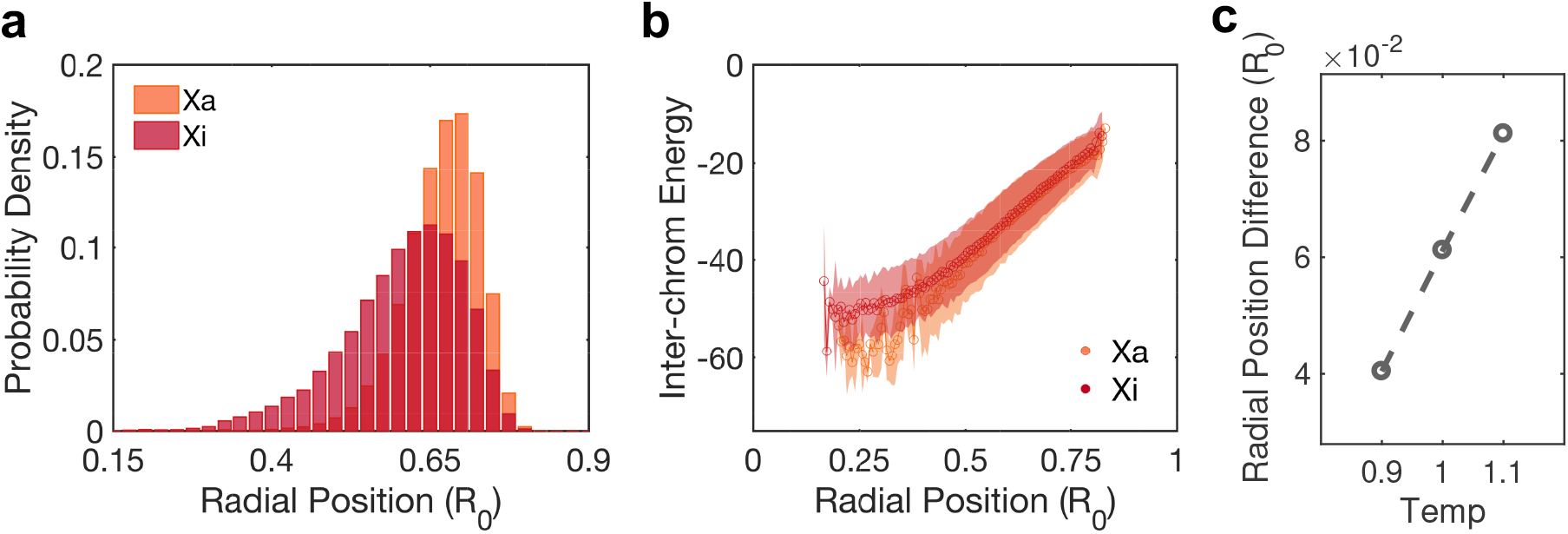
Contribution of configurational entropy to the interior localization of the inactive *X*-chromosome. (a) Probability distribution of the radial position for the active (orange) and inactive (red) *X*-chromosome. (b) Average inter-chromosomal contact energy between the *X* chromosomes and the rest of the genome as a function of chromosome radial position. The shaded areas represent the standard deviation in the contact energy estimated from the simulated structural ensemble. (c) Dependence of the radial position for the two *X* chromosomes as a function of temperature. *R*_*o*_ is the radius of the nucleus used in polymer simulations.

Its combined utility for structural reconstruction and mechanistic exploration renders the modeling framework presented here useful for studying genome organization. In the meantime, we emphasize that the framework is general and additional experimental data other than Hi-C^21,63^ can be further incorporated to improve model accuracy. It would be particularly interesting to integrate the Hi-C based model with results obtained from super-resolution imaging^55,64–67^ to characterize single-cell genome structures.

## Methods

### Data Processing

#### Hi-C Contact Maps

Data were downloaded from the Gene Expression Omnibus database using SRAtoolkit version 2.9.0 and the SRAdb R/Bioconductor package. The GM12878 data were downloaded from the series number GSE63525. Only samples generated using in situ Hi-C protocol were considered for further analysis. A catalog of phased SNPs was downloaded from the thousand genome project database for the individual NA12878.

Hi-C reads for GM12878 were mapped to the human reference genome version hg19 where heterozygous SNPs for NA12878 were masked with N’s. For reads whose alignment to the reference genome overlapped with a SNP, the read was assigned to either the maternal allele if the sequence of the read matched with the sequence of the maternal allele or to the paternal allele if the sequence of the read matched the sequence of the paternal allele. A sequence fragment was considered a valid read pair if one read mapped to a restriction fragment different to the corresponding read pair. HiC-Pro^68^ was used for both the mapping step and to generate a list of valid Hi-C contacts. In order to build allele-specific contact maps, valid contacts where at least one read overlapped with SNPs were considered.

Three genome-wide contact maps of 1 megabase resolution were generated: (1) a matrix built with contacts for which at least one read overlapped with a SNP and contained the maternal allele, a matrix built with valid contacts for which at least one read overlapped with a SNP and contained the paternal allele and (3) a matrix built with all valid contacts ignoring information of the alleles. The three raw genome-wide contact maps were then balanced using the Iterative Correction and Eigenvector decomposition (ICE) method.^69^

#### Compartment Profile

To assign compartment labels to each 1 megabase bin in the genome, the balanced haploid contact map *X* = *x*_*i j*_, with *i* rows and *j* columns, were transformed into a normalized matrix by dividing each element of the matrix by the expected interaction frequency given the distance from the diagonal *k* = *i* − *j*. The expected interaction frequency (i.e. the normalization factor) was defined as the mean of the *x*_*i j*_ values with the same value of *k*. A correlation matrix was generated by estimating the pairwise correlation coefficients between all pairs of rows of the normalized matrix. Then, an eigenvector decomposition was performed on the correlation matrix and the sign of the first eigenvector was used to assign compartment labels. Since the sign of the eigenvector is arbitrary in an eigenvector decomposition, gene expression data and GC content were used to flip the sign of the eigenvectors when necessary so that positive eigenvectors always corresponded to open (*A*) chromatin and negative eigenvectors corresponded to closed (*B*) chromatin. For the central chromosome regions with missing Hi-C data, we assigned 2Mb segments outside the boundary and up to 7Mb segments next to but within the boundary as type *C*. This assignment allows the estimation of centromere contact frequency and interaction strength using neighboring contacts. All other genomic loci with missing Hi-C data were assigned as type *N*, and no explicit interactions were included for them in the polymer model.

#### Single-Cell Data

Reconstructed single-cell genome structures at the 20kb resolution for the GM12878 cell line^62^ were directly downloaded using the Gene Expression Omnibus (GEO) accession number GSE117109. The radial position of each chromosome was then determined as the center of mass coordinate of all its genomic loci. A 46 × 46 dimensional distance map can be calculated from the radial positions to directly compare with results from our polymer simulations.

### Energy function of the whole genome model

Here we provide more detailed mathematical expressions for each term of the diploid genome model energy function (Eq. 1).

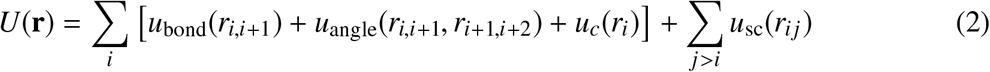

defines the connectivity of the polymer with bonding (*u*_bond_) and angular (*u*_angle_) potentials. *u*_*c*_(*r*_*i*_) corresponds to a boundary potential applied to each bead *i* to mimic the confinement effect by the nuclear envelop. The radii of the spherical boundary *R*_*o*_ was chosen to achieve a volume fraction of 0.1. *u*_sc_(*r*_*i j*_) is a soft-core potential between non-bonded pairs (*i*, *j*) formed by beads both from the same and different chromosomes. It accounts for the excluded volume effect while allowing polymer chains to cross over each other with a finite probability. Parameters used in these potentials can be found from our previous work.^38^

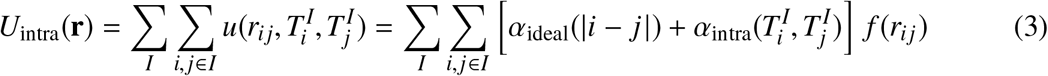

describes the type-specific intra-chromosomal potential for each chromosome *I*. We further split the potential into an ideal term, *α*_ideal_(|*i* − *j*|), that is only a function of the sequence separation between two genomic loci *i*, *j* and another term, 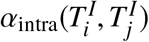, that depends explicitly on the compartment types 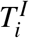 and 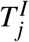. *f* (*r*_*i j*_) measures the probability of contact formation for two loci separated by a distance of *r*_*i j*_.^38^

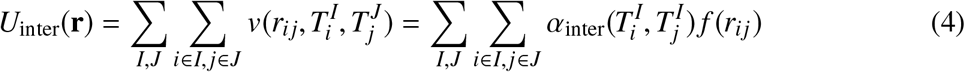

is similarly defined but for interactions between genomic loci from different chromosomes.

To capture the distinction between active and inactive *X*-chromosomes, we applied an additional potential

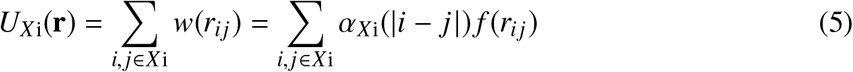

to pairs of genomic segments from the inactive *X*-chromosome. This potential is motivated by the observation that the average contact frequency as a function of genomic separation is much larger for the *X* chromosome than autosomes (Fig. S7a). Further splitting the contact frequencies into contributions from the two homologs suggests that the difference between *X* chromosome and autosomes seen in haploid Hi-C is mostly caused by condensation of the inactive *X*-chromosome (Fig. S7b). It is, therefore, reasonable to assume the active *X*-chromosome shares the same ideal potential as other autosomes and only apply the correction term to the inactive copy.

Finally, to more accurately reproduce inter-chromosomal contacts, we introduced an additional potential

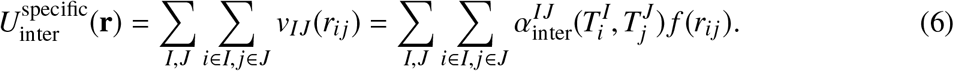

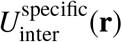 is only applied to type *A* and *B* beads. It helps to capture specific interactions encoded in the DNA sequence while retaining the simple representation of chromosomes using *A*/*B* compartments. Since haploid Hi-C data do not report contact frequency between homologous chromosomes, this potential was not defined for those pairs.

For most of the parameters introduced in Eqs. 3–6, {*α*_intra_, *α*_inter_, *α*_*Xi*_, 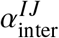}, a corresponding constraint based on haploid Hi-C data can be defined. Values of these parameters can then be fine tuned with an iterative algorithm to ensure that ensemble averages calculated using simulated genome structures match with Hi-C constraints.^27,38^ Detailed mathematical expressions for the constraints are provided in the SI.

All parameters for type *N* were set as zero due to the lack of corresponding Hi-C data. The interaction energy between centromeres was approximated with values derived for pericentromeric regions. To ensure the accuracy of this approximation, we only designated segments close to pericentromeric regions as type *C* in the polymer model (see *Data Processing: Compartment Profile*). Finally, for haploid Hi-C, the contact frequency between homologous chromosomes are not measured explicitly but grouped together with intra-chromosomal contacts. We, therefore, approximated interactions between homologous chromosomes using the energy derived from heterologous pairs (see SI for details).

### Details for molecular dynamics simulations

We carried out polymer simulations using the molecular dynamics package LAMMPS^70^ in reduced units. Simulations were maintained at a constant temperature of *T* = 1.0 via the Langevin dynamics with a damping coefficient of *γ* = 10.0 and a time step of *dt* = 0.01. The initial configuration for these simulations was generated as follows. We first placed the chromosomes consecutively on a cubic lattice with a length of 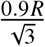, where *R* = 19.7*σ* is the radii of the spherical confinement introduced to ensure the volume fraction of the genome as 0.1. We then relaxed the genome structure with a 100,000-step-long simulation under the homopolymer potential *U*(**r**) (Eq. 2). The end configuration of this simulation was used as the starting structure for all subsequent simulations. We note, however, that all the results present in the manuscript correspond to equilibrium averages and are independent of the initial configuration.

Parameters in the energy function were determined through an iterative algorithm introduced by us.^27,38^ We started the first iteration of molecular dynamics simulations from the equilibrated configuration generated above. All subsequent simulations were initialized with the end configurations from the previous iteration. During each iteration, four independent 20-million-time-step-long simulations were carried out and genome conformations were recorded at every 2000 timesteps to calculate the ensemble averages. Parameters in Eq. 2–6 were then updated to minimize the difference between the simulated 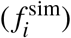 and experimental 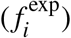 constraints whose detailed expressions are provided in the SI. The simulation error defined as 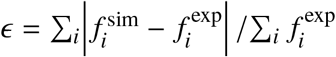 reaches a value of less 5% in the last round of iteration. With the parameterized energy function, we then performed four independent 40-million-time-step-long simulations and recorded genome conformations at every 2000 timesteps for data analysis. Similar optimization procedures were carried out for the three perturbations presented in Fig. 5.

## Acknowledgements

We thank Wenjun Xie and Xinqiang Ding for helpful discussions. This work was supported by the National Science Foundation Grant MCB-1715859 and the National Institutes of Health (Grant 1R35GM133580-01).

## Author contributions

Y.Q., A.R., S.E.J., M.J.A., B.E.B.,and B.Z. conceived the research; Y.Q., A.R. and B.Z. performed research and analyzed the data; Y.Q. and B.Z. wrote the paper with input from all other authors.

